# Theta band-power shapes amyloid-driven longitudinal EEG changes in pre-clinical Alzheimer’s Disease

**DOI:** 10.1101/2022.02.04.479130

**Authors:** Giuseppe Spinelli, Hovagim Bakardjian, Denis Schwartz, Marie-Claude Potier, Marie-Odile Habert, M. Levy, Bruno Dubois, Nathalie George, the INSIGHT-preAD study group

## Abstract

Alzheimer’s Disease (AD) includes progressive symptoms spread along a continuum of pre-clinical (pre-AD) and clinical stages. Pre-AD refers to cognitively healthy individuals with presence of positive pathophysiological biomarkers of AD (i.e., markers of amyloidopathy and tauopathy). Although numerous studies uncovered the neuro-cognitive changes of AD, very little is known on the natural history of brain lesions and modifications of brain networks of pre-AD. To address this issue, we analysed resting-state EEG data of 318 cognitively healthy individuals with subjective memory complains from the INSIGHT-preAD cohort at the time of their first visit (M0) and two-years later (M24). Using 18F-florbetapir PET-scanner, subjects were stratified between amyloid positive (A-; n=230) and amyloid negative (A+; n=88) groups. Differences between A+ and A- individuals were estimated at source level in each band of the EEG power spectrum. At M0, we found an increase of theta-band power in the mid-frontal cortex in A+ compared to A-. No significant association was found between mid-frontal theta power and the individuals’ cognitive performance. While the very same effect was not replicated at M24, theta-band power increased in A+ relative to A- individuals in the posterior cingulate cortex and the pre-cuneus. Furthermore, alpha band revealed a peculiar decremental trend in posterior brain regions in the A+ relative to the A- group only at M24. These results provide the first source-level longitudinal evidence on the impact of brain amyloidosis on the EEG dynamics of a large-scale, monocentric cohort of pre-AD. Theta-band power increase over the mid-frontal and mid-posterior cortices suggests an hypoactivation of the default-mode network in individuals at-risk of AD and a non-linear longitudinal progression of the AD-spectrum.

## 1. Introduction

Alzheimer’s Disease (AD) is the most prevalent neurodegenerative disease and the most common form of dementia (approximately 60-80% of dementia cases; Alzheimer’s Association, 2015). AD primarily affects elderly individuals (∼65 years; Alzheimer’s Association, 2015) and includes cognitive symptoms and biological signs spread along a continuum of pre-clinical (pre-AD) and clinical stages. Symptoms mainly include impairments in cognitive functions and loss of memory, while biological signs consist in potentially irreversible neuro-physio-pathological alterations including senile plaques, neurofibrillary tangles and inflammatory changes (Braak and Braak, 1991; Duyckaert et al., 2009). Despite many scientific efforts, the large heterogeneity of AD phenotype and the non-linear progression of both signs and symptoms render clinical and pharmacological trials mostly ineffective. Indeed, no efficient drug exist to slow the progression of AD (success rate for approval is ∼0.4%; Cummings et al., 2014). Cognitive therapies limit to link life-long cognitive stimulations and trainings to an extended tolerance to the onset of AD symptoms (i.e., the so-called “cognitive reserve” hypothesis; Stern, 2002; 2012) or other forms of cognitive impairment (Olazaran et al., 2004; Solé-Padullés et al., 2009; Amodio et al., 2017), but no decisive solutions has emerged yet. Therefore, AD has become a huge public health problem, counting a broad increasingly number of people living with dementia worldwide (∼46.8 million individuals, predicted to double every 20 years; Prince et al., 2015).

From a pathophysiological point of view, the most credited theory on the AD pathogeneses is the amyloid cascade hypothesis (Hardy and Higgins, 1992; Selkoe et al., 2016) which posits that brain β-amyloid (Aβ) accumulation is the key event which triggers neuronal damage, synaptic function impairments and ultimate widespread neurodegeneration. Crucially, all these events appear many years before the onset of clinical symptoms (Walsh et al., 2017; Jack et al., 2013) and are thought to cause a so-called disconnection syndrome (Brier et al., 2014): brain functional and structural connections become progressively and irreversibly disrupted. Moreover, Aβ load is supposed to behave as an accelerator of other consequent biochemical events (Williams, 2006) together with genetics (Chiesa et al., 2018) and sex (Chiesa et al., 2019; Dumitrescu et al., 2019; Cavedo et al., 2018; see Ferretti et al., 2018 for a review), thus resulting in additive risks of developing AD. Therefore, two phases are now considered in the AD continuum: a preclinical phase of more than 15 years and a clinical one which encompasses a prodromal (predementia) and a dementia stage (Dubois et al., 2016). Given the lack of efficacy of disease-modifier therapies in patients with dementia, scientific interest has grown on preclinical stages of AD (pre-AD), i.e. the stage in which the earliest neuropathological events appear (i.e., amyloid- or tau-pathology) in the absence of AD-related cognitive symptoms (Dubois et al., 2016). Characterizing the neurophysiological patterns of pre-symptomatic individuals with high β-amyloid load has thus become one of the most crucial challenges in clinical neuroscience, with the hope of being able to predict the progression to AD.

In this vein, electroencephalography (EEG) has been used for many years to acquire a direct, non-invasive view of human brain activity in condition of physiological and pathological aging (i.e., AD or other dementias). Indeed, EEG allows an extremely precise discrimination of the temporal hierarchy governing distributed brain networks at different frequencies, i.e., 1–4 Hz (delta), 4–8 Hz (theta), 8–13 Hz (alpha), 13–30 Hz (beta), and >30 Hz (gamma), thus allowing to investigate the information that these canonical frequencies may convey on various mental and physiological states. In this regard, physiological aging is characterized by a marked “slowing” of both the background EEG (see Klimesch, 1999, for a review) and a decreased power of the occipital alpha-band which, in turn, is typically accompanied by an increase of slower frequencies, i.e., delta and theta (Babiloni et al., 2006). Despite this, healthy seniors often show plastic compensatory brain mechanisms which contribute to relatively long-term functional maintenance and absence of any symptoms. When symptoms become sufficiently evident, albeit not severe enough to exceed standard critical criteria for AD, Mild-Cognitive Impairment (MCI) emerges. A meta-analysis conducted on 56 non-demented seniors, 106 MCI, and 108 AD patients showed an intermediate magnitude of alpha power in parieto-occipital brain areas in MCI with respect to the other two groups (Babiloni et al., 2011). Such a decrease of alpha power has been linked to a dys-regulation of the thalamo-cortical and cortico-cortical brain networks governed by the cholinergic brainstem pathway (Dringenberg, 2000), which is thought to be responsible of the transmission of sensori-motor information and the retrieval of semantic information (Klimesch, 2012). Besides alpha, lower frequencies of the EEG power spectrum have also been associated to different neural peculiarities of MCI. More specifically, while delta power has been inversely associated to cortical atrophy (Babiloni et al., 2006), both theta band-power and theta interhemispheric coherence were found to be predictive of the decline from MCI to AD (Rossini et al., 2006). Taken together, these findings corroborate the idea that MCI can be considered as a transitive stage between physiological aging and AD, not only from a clinical (Misra et al., 2009), but also from an electrophysiological point of view. For what concerns AD, EEG studies convincingly showed an increase widespread delta and theta power and a decreased posterior alpha power in comparison with both normal seniors and MCI (Penttilä et al., 1985; Huang et al., 2000; Moretti et al., 2004; Musaeus et al., 2018; see Babiloni et al., 2012 for a review). Interestingly, alpha decrease is generally associated to impaired cognitive function as indexed by neuropsychological tests and batteries, such as the Mini-Mental-State Examination (Jeong, 2004). Characterisation of posterior alpha power has also allowed discrimination among different phenotypes of dementias (Del Percio et al., 2018; Durongbhan et al., 2019). Decreased of posterior alpha power or of the so-called individual-alpha peak (IAF; i.e., the frequency associated to the strongest EEG power within the standard alpha range) differentiates mid-AD from early-AD (Gallego-Jutglà et al., 2015), cerebrovascular dementia, fronto-temporal dementia, and early-stages of AD (Moretti et al., 2004; Al-Qazzaz et al., 2018). Moreover, classification of theta power has been used to explain subtle working memory changes in AD (Goodman et al., 2018), as well as to differentiate AD dementia from Lewy-body dementia (Peraza et al., 2018). Relevant for the present work are studies showing the association between additional biological parameters and cortical EEG markers in AD. Specifically, increased genetic risk of AD - as indexed by presence of *Apolipoprotein-E* (*APOE*) ε4 genotype – is thought to affect spontaneous electrical activity (Cuesta et al., 2015), in both theta and beta bands (Lehtovirta et al., 1996), as well as global EEG connectivity (Gonzales-Escamilla et al., 2015; Michels et al., 2017; Koelewijn et al., 2019). In this respect, although connectivity measures have become fine-grain markers to better characterize changes of brain network dynamics, findings in AD are not homogeneous and there is still no consensus on which connectivity analysis (e.g., local connectivity, global connectivity, network assortativity, or graph analysis) or which measure of connectivity (e.g., real/imaginary coherence, phase-locking value, network-node, or entropy) may reliably reflect brain changes along the AD continuum (Stam et al., 2002; Brunovsky et al., 2003; Stam et al., 2005; Babiloni et al., 2009; Yu et al., 2016; de Haan et al., 2017; Guillon et al., 2017; Babiloni et al., 2018; Vecchio et al., 2018).

In view of all this evidence, investigating the EEG changes of pre-clinical AD has become tremendously relevant for the scope of AD research. To the best of our knowledge, only a few EEG studies have been reported in pre-clinical AD. Target cohorts typically include individuals with increased risk to convert to AD, such as subjective cognitive/memory complainers with and/or without positive biomarker status for AD, e.g., cortical amyloidosis (Dubois et al., 2016; Jonker et al., 2000). In particular, Babiloni et al. (2010) revealed an increase of frontal lower frequency bands at source level in a group of 53 subjective memory complainers as compared to 79 healthy seniors and 143 MCI (both amnesic and not-amnesic). Prichep et al. (2006) reported that mid-frontal theta band power was able to predict the conversion from non-pathological aging to MCI in a group with subjective cognitive decline. This finding was replicated by Gouw et al. (2017) who showed that theta power increase on mid-frontal scalp locations characterized the conversion to MCI in a sub-group of non-demented seniors with an amyloid positive biomarker status. Noteworthy, the scarce control of i) any underlying neuro-pathology (i.e., amyloid status or genetic profile), ii) the inter-subject variability due to scarce homogenisation of the studied population, iii) the exclusion of neurological comorbidities and mixed-type symptoms, and the relatively small sample sizes, limit the generalisation of these findings.

The ongoing longitudinal INSIGHT-preAD cohort (INveStIGation of AlzHeimer’s PredicTors in subjective memory complainers) was setup to address these issues (Dubois et al., 2018). First resting-state EEG investigations on this cohort at the baseline period, that is, at the time of the first visit (i.e., M0), revealed contradictory results. While Teipel et al. (2018) revealed no association between amyloid burden and scalp-level regional EEG connectivity, Gaubert et al. (2019) mainly showed an increased connectivity within a fronto-central scalp network as a function of the amyloid status and the degree of neurodegeneration.

In the present work, we capitalise on these findings to further evaluate the impact of cortical amyloid load on the resting-state EEG pattern of the INSIGHT-preAD cohort by means of longitudinal, source-level EEG analyses. Source-level analysis guarantees a more fine-grained exploration of the neural dynamics of this population, whilst longitudinal approach provides with a more robust test of any long-term effect as well as a characterisation of the progression of the disease. In addition, we tested for genetic influence (presence of the *APOE* ε4 allele), for the confounding effects of sex and age, and for any association between EEG features and individuals’ cognitive performance.

## 2. Methods

### 2.1 Sample

Participants were selected from the INSIGHT-preAD study (Dubois et al., 2018), a French monocentric cohort including longitudinal data of 318 seniors without objective cognitive impairment (185 females; mean-age: 76.1 years, range: 70-85 years) recruited from the Institute for Memory and Alzheimer’s Disease (IM2A) at the Pitié-Salpetriere Hospital of Paris. All of them had complained about memory issues for more than 6 months before the recruitment. To be included in the cohort, participants should have normal cognitive abilities (Mini-Mental State Examination score ≥ 27; Clinical Dementia Rating score = 0) and no objective memory deficits (Total recall score ≥ 41 at the Free and Cued Selective Reminding Test). Written informed consent was collected at the time of the first visit (i.e., baseline or M0). The study conformed to the 1975 Declaration of Helsinki and was approved by the local ethics review board. To study longitudinally the neurophysiological profile of the cohort, data referring to the first (M0: 318 participants) and to the 2^nd^ year visit (M24: 279 participants) were extracted. Following quality-check and artefact rejection (see below), the data of 272 subjects, matched across the two visits (M0, M24) were included in the analyses (see Table 1 for further details).

**Table 1.** Demographic characteristics, global cognitive profile, amyloid status, and APOE genotype of the sample. SD: standard deviation.

|  |  | **M0** | **M24** |
| --- | --- | --- | --- |
| **Participants** (female) |  | 318 (185) | 279 (172) |
| **Education** (in years) |  | 6.2 ± 2.1 |  |
| **Age ± SD** |  | 76.1 ± 3.5 |  |
| **MMSE ± SD** |  | 28.7 ± 0.96 |  |
| **FAB ± SD** |  | 16.4 ± 1.73 |  |
| **FCSRT-TR ± SD** |  | 46.1 ± 1.98 |  |
| **Amyloid**<br>**status** | **A- \| APOE ε4+** | 230 / 25 | 206 / 24 |
|  | **A+ \| APOE ε4+** | 88 / 33 | 75 / 29 |

### 2.2 Neuropsychological Assessment

Patients’ cognitive profile was assessed by a comprehensive neuropsychological battery including: the MMSE (Derouesne et al., 1999), the Clinical Dementia Rating (CDR, Morris, 1993), the Digit Span (Wechsler, 1997), the Free and Cued Selective Reminding Test (FCSRT; Buschke, 2002), Letter and Category Verbal Fluency test (Beton, 1986), the Rey Complex Figure Copy (Fastenau et al., 1999), the Trail Making Test (Tombaugh, 2004), the Frontal Assessment Battery (FAB, Dubois et al., 2000), the Memory Capacity Test (MCT, Rentz et al., 2013) and the Digit Symbol Substitution Test (DSST, McLeod et al., 1982). Further details are provided in Table 1.

### 2.3 PET-amyloid scan

A Philips Gemini GXL CR-PET scanner served to measure the amyloid uptake in the brain. Scans were acquired 50 (±5) minutes after injection of ∼370 MBq (range: 333-408 MBq) of 18F-florbetapir. The acquisition consisted in 3×5 minutes frames, 128×128 acquisition matrix with a 2×2×2 mm^3^ voxel size. Images were reconstructed by means of the LOR-RAMPLA algorithm with 10 iterations and a smooth post-reconstruction filter was applied. Frames were then realigned, averaged and quality-checked by the CATI team (http://cati-neuroimaging.com). Composite cortical ROIs (left/right-Pcu, PCC and ACC, parietal, temporal and orbitofrontal cortices) were derived and a reference region (i.e., pons and whole cerebellum) was placed in the individual native PET space. Parametric PET images were created by dividing each voxel to the mean activity of the reference region. Standard amyloid uptake value ratios (SUVr) were extracted by averaging the mean values across all cortical ROIs. Based on a linear conversion of the CAEN method to the INSIGHT cohort (Besson et al., 2015; Habert et al., 2018), individuals with a SUVr > 0.79 were classified as amyloid positive (A+) while individuals with a SUVr ≤ 0.79 were classified as amyloid negative (A-) (see Table 1; Dubois et al., 2016; Habert et al., 2018).

### 2.4 APOE genotype

DNA was extracted from frozen blood samples by applying the 5Prime Archive Pure DNA purification method. Genotypes were determined using Sanger method (see Dubois et al., 2018 for details). Individuals were then split in groups based on their *APOE* genotype and assigned to sub-samples of carriers (ε4+) or non-carriers (ε4-) of at least one *APOE* ε4 allele (i.e., ε4/ε4 and ε4/ε3) (Table 1).

### 2.5 MRI recording

A 3T Siemens TRIO MRI scanner (Siemens Medical Solution, Erlangen, Germany) equipped with a 12-channel head coil for signal reception was used for MRI acquisition. For the anatomical study, 3D TurboFlash sequences were performed (sagittal orientation, 2300 repetition time, 2.98 ms echo time, 900 inversion time, 9° flip angle, 176 slices with a 1 mm thickness, field of view of 256*240 mm, and a bandwidth of 240 Hz/Px.

### 2.6 EEG recording and analysis

Resting-state EEG (rsEEG) was acquired at 250 Hz using a 256-channel whole-head cap (EEG System GES 300, Electrical Geodesic, Inc) and amplified by the EGI NetAmp300. No modification of the signal was applied at recording. The reference electrode was Cz. Electrodes impedance was kept below 50 kΩ. Each rsEEG was acquired in two separate 2-minutes runs. Each run consisted in two consecutive sequences of 60 seconds recording in which participants were instructed to keep their eyes closed (EC; 30 seconds) and opened (EO; 30 seconds). A low- or high-frequency tone was delivered to indicate the onset of each EC or EO condition, respectively. To minimize blinking, each rsEEG run was preceded by a fixation cross (2 seconds) presented at the center of a computer-screen against a black background. The NetStation 4.4 software (EGI) was used for signal acquisition, and the E-Prime software (Psychology Software Tools, Inc, Pennsylvania, USA) allowed the synchronization of marker-events (onset and end of each EC / EO periods) in each recording sequence.

Neural time series were pre-processed using the Fieldtrip toolbox (release: July 2017; Oostenveld, 2010) in Matlab r2016a (The MathWork, Inc). Each dataset was filtered with a 1-80 Hz band-pass FIR filter (one-pass, zero-phase). Artifact detection was restricted to scalp-electrodes (n =173) and carried out by means of a thresholding procedure, such that three metrics were calculated for each sensor along the whole time-series (i.e., variance, amplitude-range and minimum voltage value) and then transformed in z-scores based on the mean and the standard deviation of the whole scalp-electrodes set. Sensors with |z-scores| ≥ 2.5 for at least one of the three metrics, were marked as artifactual and excluded from the analysis. On average, 5.4 channels (range: 1-13) were rejected for each participant. The resulting artifact-free channels were used to re-reference the time series to a common-average referencing. A visual inspection of the continuous data was carried out to exclude segments of data exhibiting any residual aberrant activity (mainly due to muscular artifacts). Then, an Independent Component Analysis (ICA) was run to correct blinks/oculomotor artifacts. ICs showing highest correlations with signals recorded from both the peri-orbital and peri-canthal plane were removed from the original signals. On average, 2.3 ICs referring to eye artifacts (range: 1-6) were removed.

Source localization of oscillatory brain activity was carried out by means of the beamformer method. For each subject, EC and EO EEG signals were band-pass filtered (FIR filter; one-pass, zero phase) into 7 frequency bands (delta [2-4 Hz], theta [4-8 Hz], alpha_1_ [8-10 Hz], alpha_2_ [10-13 Hz], beta_1_ [13-20 Hz], beta_2_ [20-30 Hz], and gamma [30-40 Hz]). For each of these band-pass filtered traces, the data covariance matrix was calculated after merging EO with EC. A common-spatial filter was obtained by means of a Linearly Constrained Minimum Variance (i.e., LCMV) beamformer and multiplied to each EC and EO condition to project the data in the source-space. The forward solution (Boundary Element Method) was obtained from a dipolar model based on a 3-compartments mesh (i.e., brain, skull, scalp) created on each individual’s MRI through FreeSurfer processing as implemented in the FieldTrip toolbox. Voxel positions were derived from individual’s MRI and normalized to a 5-mm grid covering the entire brain. Alignment between the MRI and the EEG coordinate system was carried out by marking three fiducial points (i.e., nasion, left- and right- pre-auricular) on individual MRIs. Source-level data were then normalized in the MNI space.

Frequency analysis was performed at voxel-level by means of a multi-tapered Fast Fourier Transform (FFT) using a Hanning taper. Band-power was computed by taking the squared magnitude of the real and imaginary Fourier spectra. The relative power was obtained by dividing the power in a single band by an estimate of the summed power, i.e. the sum of the power in the 7 bands. These values were ultimately transformed into z-scores (using the mean and the standard deviation across voxels) to enable comparisons between subjects and frequencies. To exclude outliers from each visit, a 95% confidence interval was computed across subjects using the global power over all the frequency bands, such that subjects exhibiting values > 95% were discarded from the analysis.

To test statistically the effect of brain amyloidosis on source-level power for each visit a Monte Carlo cluster-based permutation approach was implemented (Maris et al., 2007), with 5000 iterations. For each frequency band, a permutation distribution of the significance probabilities for independent-samples t-tests between A- and A+ was calculated. The significant threshold of both cluster-alpha and alpha was set at .05, and the maximum of the sum of t-values within each cluster was considered as the test statistic (Maris and Oostenveldt, 2007). To circumvent the issue of unequal variance and sample-size of the two groups (i.e., A+ and A-), a bootstrap-like procedure was applied to the permutation test, as described in Mewhort et al. (2009). Data from the larger group (i.e., A-) were selected at random to create a sample of equal size as the smaller group (i.e., A+) and then the Monte-Carlo cluster-based permutation test was run on this subset of the data. This procedure was implemented 100 times, each time noting whether and where (in which brain region) the independent-samples t-test rendered any significant cluster (positive and/or negative). The proportion of cases (out of 100) in which a given significant effect replicated was calculated to estimate the reliability of the initial cluster-based test.

A further exploration was carried out on the link between rsEEG activity and the severity of brain amyloidosis. Band-power data were extracted from the significant cluster of voxels and related to SUVr values distributed in 7 evenly spaced cumulative probabilities quantiles in order to have an equal number of cases per probabilities quantile. Then, a between-subjects ANOVA was run using the SUVr as independent variable (7 levels). Bonferroni method was used to correct for multiple comparisons.

The association between subjects’ electrophysiological profile and cognitive abilities was assessed by means of a linear regression model predicting regional specific band-power spectra from individuals’ memory performance (as indexed by the FCSRT-TR test).

To further explore the additive contribution of the APOE status, a between-subjects ANOVA was run, considering the power values extracted from the significant cluster(s) as the dependent measure, the amyloid status (2 levels: A- and A+) and the APOE status (2 levels: ε4+ and ε4-) as independent variables, and both the sex (dichotomic) and the age (continuous) as covariates.

## 3. Results

### 3.1 Cortical dynamics at M0

#### Electro-cortical markers of brain amyloidosis

The contrast between the neurophysiological profile of A- and A+ subjects revealed significant differences in the theta-band [4-8 Hz] only, at the time of the first visit (i.e., M0; Figure 1a and 1b). A+ exhibited an increase of theta-band power in comparison with A- (cluster-statistic: -98.33, t= 5.01, p < 0.02 [cluster-corrected]; Figure 1c). This effect localized on a bilateral mid-frontal cluster of voxels including the Rectus gyrus (Rec), the frontal Superior (fsOrb) and Medial (fmOrb) Orbital cortex and the Anterior Cingulate Cortex (ACC). The reliability of the test, i.e. the number of times in which the test resulted in the same identical effect, was 82% (82 cases out of 100). Moreover, the vincentization of amyloid load showed that the theta enhancement was parametrically modulated by the severity of brain amyloidosis (F_(6, 265)_ = 6.3, p < 0.001, η^2^ = 0.99): subjects with a higher accumulation of cortical amyloid (i.e. quantiles 6-7) showed a greater increase of mid-frontal theta-power in comparison of those with a below-threshold amyloid deposition (i.e., quantiles 1-5; p < 0.01; Figure 1d). This pattern was replicated by a Linear Regression Model (r^2^ = 0.08, p < 0.001) considering continuous amyloid values as predictors of mid-frontal theta power (Figure 1e).

**Figure 1.**
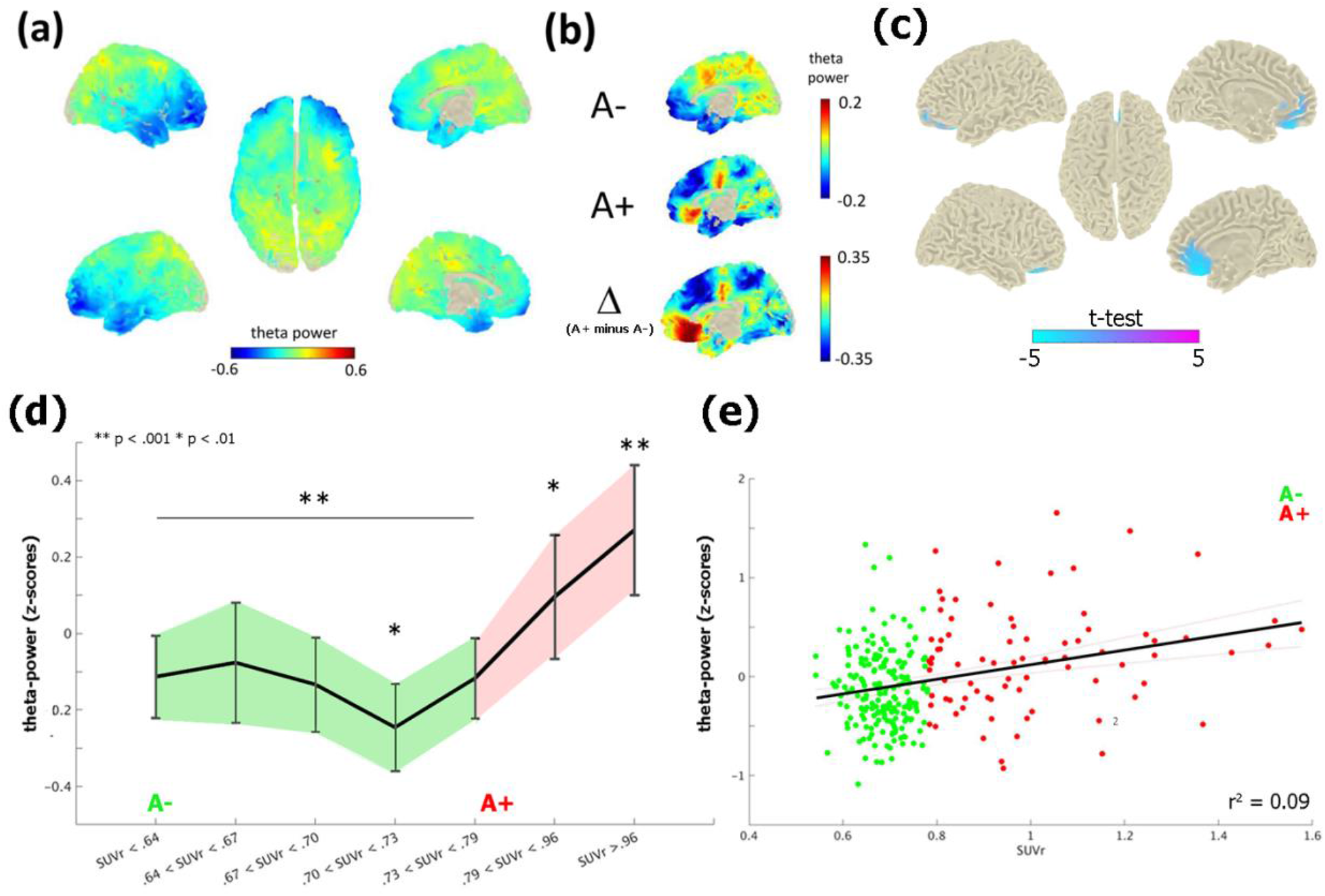
Theta-band power distribution on the brain surface at the time of the first visit (i.e., M0). Panel (a) depicts the grand-mean across the whole sample. Panel (b) shows the difference between A+ and A-. Values are expressed in z-scores. Panel (c) highlights the contrast between the groups (i.e., A- vs A+) computed through an independent-samples t-test. Significant probabilities are corrected at cluster-level by means of a Monte Carlo permutation approach. Note: negative t-test (in cyan) on the mid-frontal brain regions reflects ampler theta-power for A+ relative to A-. Panel (d) depicts the significant increase of theta-power as a function of amyloid deposition. Continuous values of amyloid are distributed in 7 quantiles, each including averaged theta-power values of ∼38 subjects. In this, individuals with highest amyloid deposition (i.e., quantile 7; A+) shows a higher (** = p < 0.001) increase of mid-frontal theta-band power with respect to A- (i.e., quantiles 1-5). A moderate effect (* = p < 0.01) is also shown between the middle (i.e., quantile 4) and medium-high stages of amyloid deposition. Vertical bars indicate 95% confidence interval. The scatterplot in panel (e) shows the linear regression between continuous values of amyloid and mid-frontal theta power.

#### Exploring any additive effect of APOE genotype, sex, and age

The significant main effect of amyloid load on mid-frontal theta power (F_(1, 271)_ = 19,34, p < 0.001) was not affected by the confounding covariates of age (p = 0.52) and sex (p = 0.99), and - importantly – there was neither any main effect of *APOE* genotype (p = 0.75) nor any interaction between the amyloid (i.e., A- vs. A+) and the *APOE* status (ε4- vs. ε4+; p = 0.64; Figure 2c).

**Figure 2.**
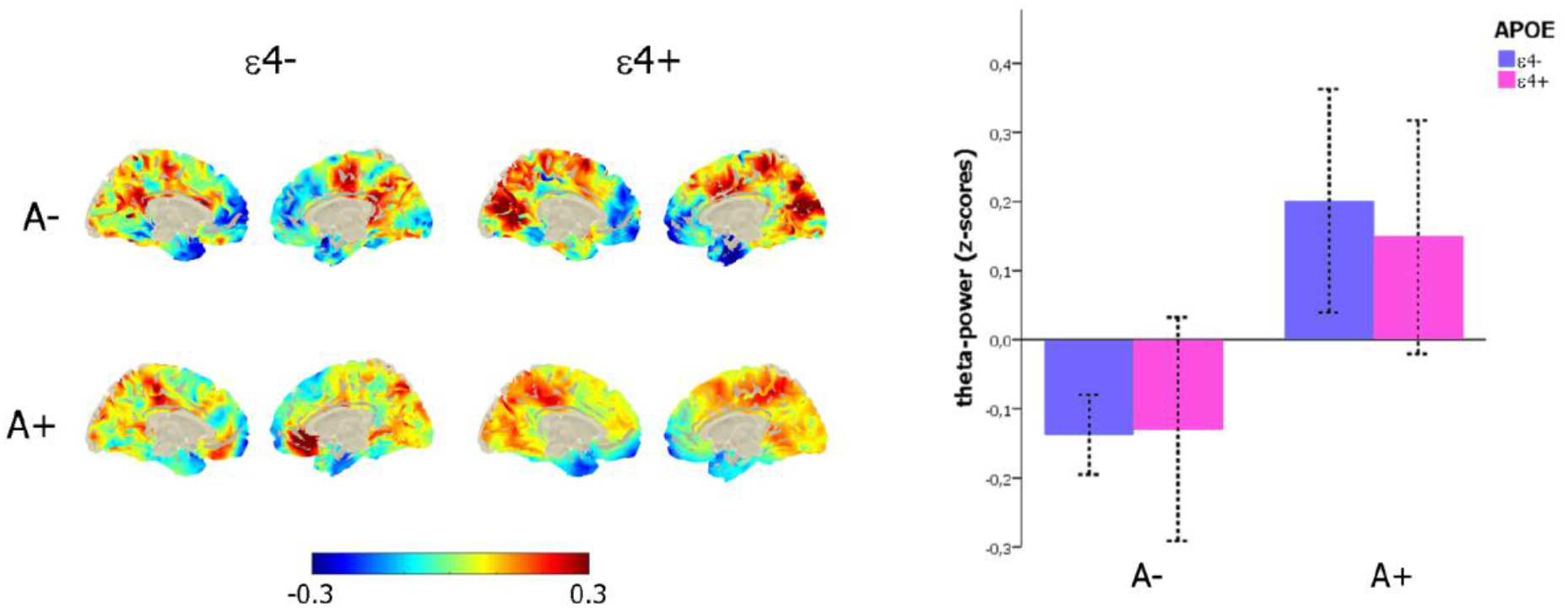
Link between mid-frontal theta-band power, amyloid deposition and APOE genotype. Left-column depicts the theta-power distribution on the brain surface (left-column) as a function of the amyloid (i.e., A- vs. A+) and APOE (i.e., e4- vs. e4+) status. The bar-plot (right-column) represents the variation of mid-frontal theta power as a function of amyloid (i.e., A- vs. A+) and APOE (i.e., e4- vs. e4+) status.

#### Linking neurophysiological patterns and cognitive performance

The regression between FCSRT-TR and mid-frontal theta power showed a non-significant inverse trend (F_(1, 270)_ = 1.67, p = 19, r^2^ = .006, r^2^_adjusted_ = .002) explained by the fact that individuals with decreased memory performance tended to exhibit higher values of mid-frontal theta power (Figure 3a). The same analysis conducted in the two groups (i.e., A+ and A-) separately, indicated that the aforementioned trend was mainly accounted for by people with a lower cortical amyloid load (i.e., A-: F_(1,198)_ = 1.92, p = .15; A+: F_(1,70)_ = 0.19, p = 0.67; Figure 3b).

**Figure 3.**
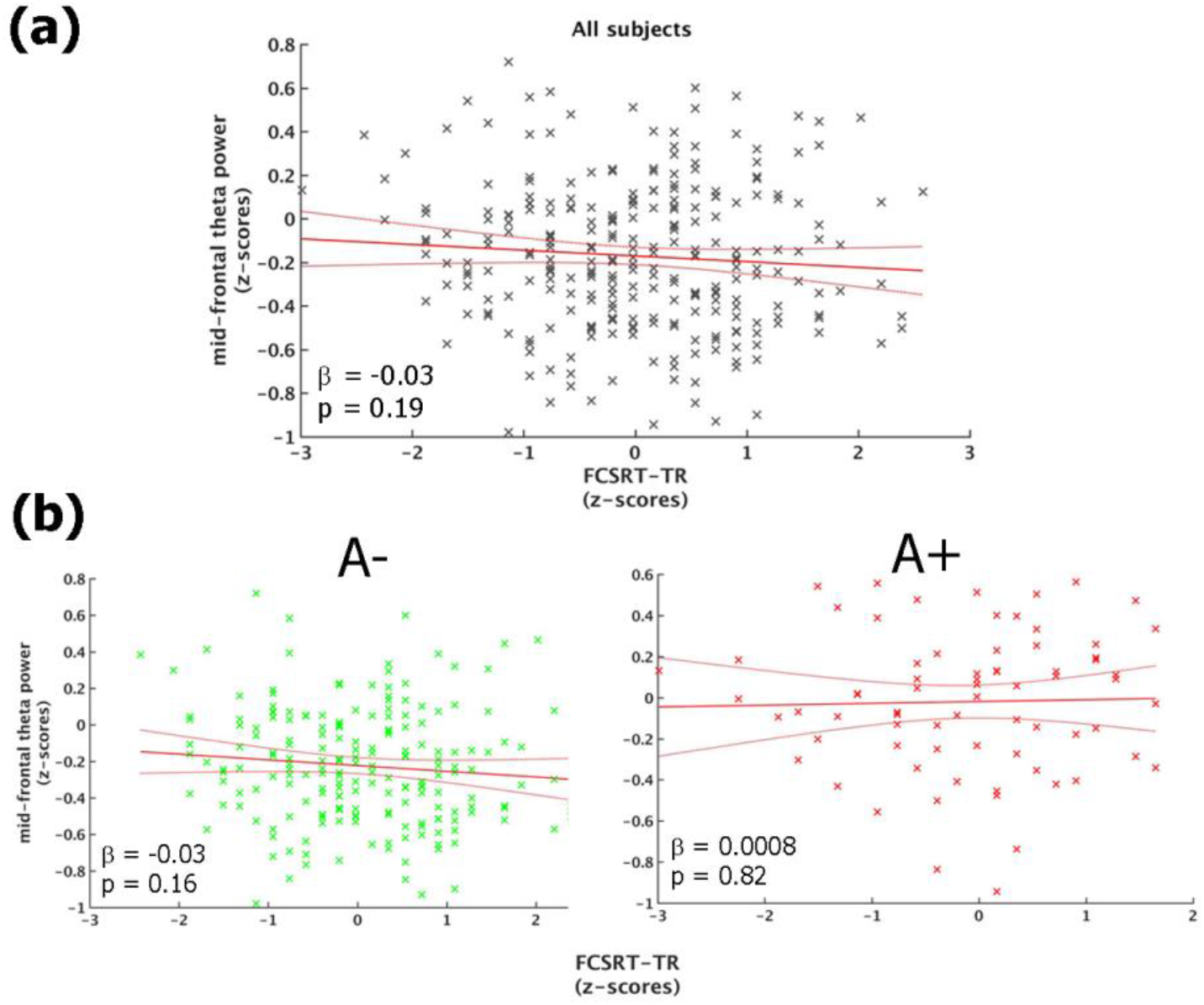
Link between mid-frontal theta-band power and memory performance (FCSRT-TR). Panel (a) shows the linear regression model carried out on the whole sample. Panel (b) depicts the same model performed on A- (left-column) and A+ (right-column), separately. All the plot shows the raw data (black/green/red stars), the fit of the model (red straight-line) with 95% confidence interval (red curved-line), its slope (β-coefficients) and the associated p-value.

### 3.2 Cortical dynamics at M24

We conducted similar analyses at the time of the second-year follow-up (i.e., M24), in all frequency-bands. The only significant effect or trend were found in the theta and alpha bands.

#### Theta band-power

The previous mid-frontal theta-band power increase shown at M0 by A+ participants was not replicated. However, the analysis showed a significant bilateral increase of theta-band power in a posterior midline brain region including the pre-cuneus (pCu), the Posterior-Cingulate cortex (PCC), and the Calcarine fissure (Calc) (cluster-statistic: -100.15, t = 4.8, p < 0.02 [cluster-corrected]; Figure 4). The bootstrap-like procedure indicated that this effect was replicated in 44% of cases when considering a random subsample of A- participants of equal size as the A+ group, pointing to a likely greater variability in the A- group.

**Figure 4.**
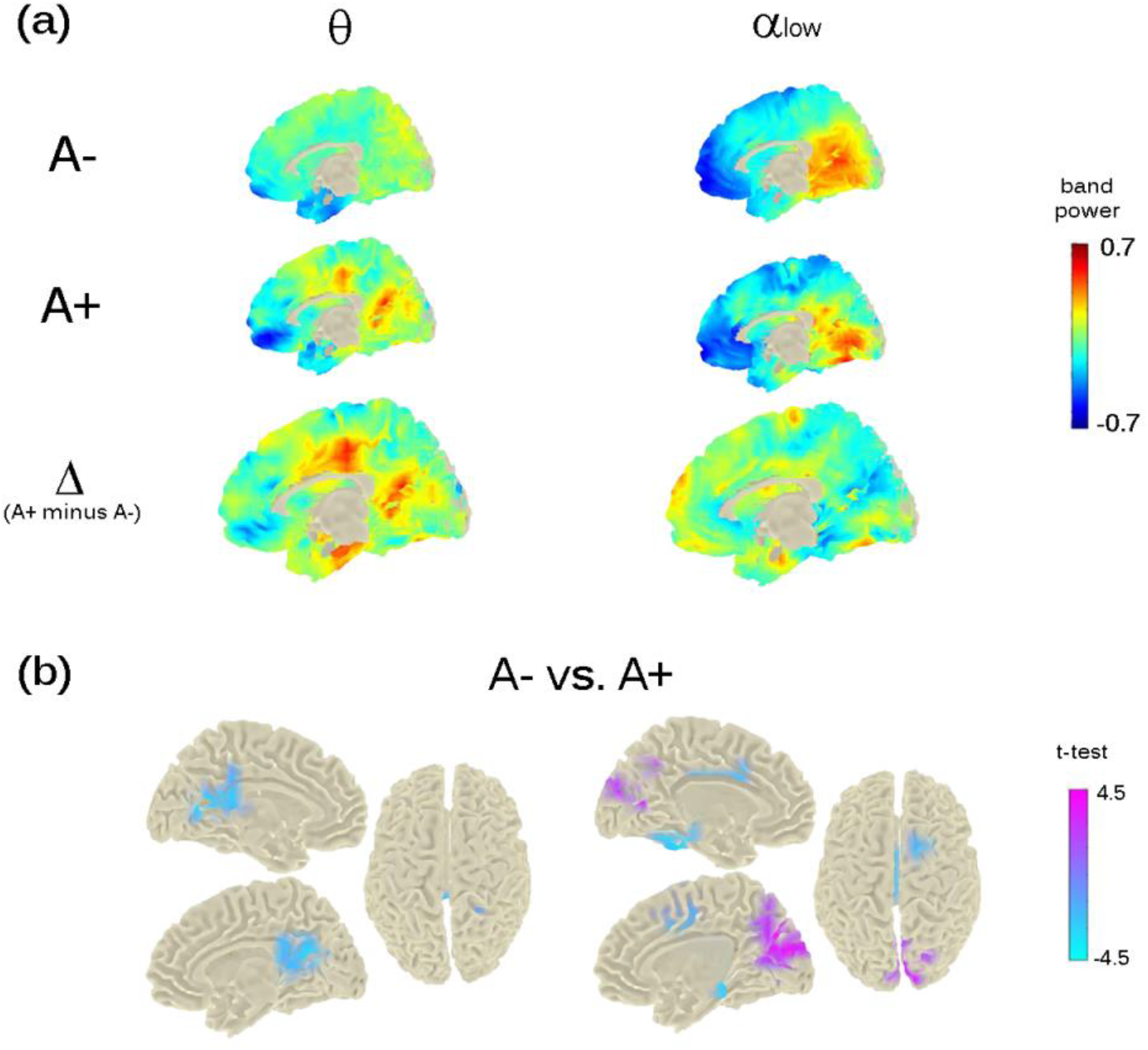
Theta- and Alpha-low band power distribution on the brain surface at the time of the second visit (i.e., M24). Panel (a) depicts the grand-mean across the whole sample for A- and A+ as well as the difference between A+ and A-. Power-values are expressed in z-scored as for M0. Panel (b) shows the independent-samples t-test computed between A- and A+. Negative t-test (in cyan) reflects ampler band-power for A+ with respect to A-; positive t-test (in magenta) reflects lower band-power for A+ with respect to A-. Importantly, significant probabilities are cluster-corrected for theta-band (left-column) and not at all corrected for alpha-low (right-column).

#### Alpha band-power

The analysis run on alpha band - considering both low-alpha and high-alpha bands – showed a significant effect only in the alpha_1_ band (8-10 Hz; p < 0.05). Individuals with an increased risk to develop AD (i.e., A+ individuals) showed a decreased power in the cortical region around the parieto-occipital sulcus and the calcarine fissure (Figure 4), in keeping with previous investigations on MCI and AD. However, this effect did not resist the cluster correction (Figure 4, top- and bottom- right raw). No significant differences were found for alpha_2_ (10-13 Hz).

#### Teasing apart the role of APOE genotype, sex, and age

The significant main effect of amyloid deposition on mid-caudal theta power at M24 (F_(1, 271)_ = 6.8, p < 0.05) was not affected by the confounding covariates of age (p = 0.12) and sex (p = 0.17). No main effect of *APOE* genotype (p = 0.58) nor any interaction between the amyloid and the *APOE* status was found. The regression between mid-caudal theta power and FCSRT-TR showed no significant effect.

## 4. Discussion

In this study, we provide the first EEG evidence of the impact of cortical amyloid deposition on the longitudinal resting-state neuro-dynamics of elderly subjective memory complainers from the standardized, large-scale INSIGHT-preAD cohort. We found an increase of theta-band power in the mid-frontal cortex in the group of individuals with a higher (A+) compared to a lower (A-) amyloid deposition at the time of the first visit (M0), regardless of the individual *APOE* status, sex, and age. No significant association was found between mid-frontal theta power and individuals’ cognitive performance. Noteworthy, while the very same effect was not replicated at the second-year follow-up (M24), theta-band power was found to be increased in A+ relative to A- individuals in the posterior cingulate cortex and the pre-cuneus. Moreover, an *ad-hoc* analysis computed on the alpha band revealed a trend to a decrease of low-alpha band power in the brain regions surrounding the calcarine cortex and the parieto-occipital sulcus, in the A+ relative to the A- group. However, this latter effect did not resist the multiple comparisons correction.

Theta neurodynamic changes have been frequently reported in previous EEG investigations on the AD continuum. In particular, widespread theta (and delta) band-power increase is one of the most prominent markers of the so-called “EEG slowing” occurring in physiological aging and has been linked to individuals’ cognitive and memory decline (Klimesch, 1999). Studies on MCI and AD further support this idea. In fact, both theta band-power increase and theta-driven interhemispheric coherence are predictive hallmarks of the conversion from MCI to AD (Rossini et al., 2006), differentiate AD from other dementias (Peraza et al., 2018; Musaeus et al., 2018). As regards the pre-AD stages, mid-frontal theta power is thought to be linked to the progression from physiological aging to the prodromal (predementia) stage of clinical AD both in the absence (Prichep et al., 2006) and in the presence (Gouw et al., 2017) of cortical amyloid deposition. A recent cross-sectional investigation on the INSIGHT pre-AD cohort also reported a significant sensitivity of mid-line theta scalp connectivity to brain amyloid load (Gaubert et al., 2019). Our data nicely fit with all these findings and bring new insights not only on the cortical sources of theta band, but also on the prominent impact that amyloid load has on these EEG sources. Such a peculiar sensitivity of mid-frontal theta rhythms to brain amyloidosis in healthy subjects at risk for AD may first be viewed in the face of the impact that early amyloid deposition (and its spreading trajectories) has on the temporo-frontal neural networks. Amyloid plaques initially deposit on the temporal neocortex and disrupt the neural circuits connecting this region with the pre-frontal cortex (Walsh et al., 2017). It has been shown that theta oscillations orchestrate the communication between pre-frontal areas and the hippocampus (Mitchell et al., 2008; Siapas et al., 2005), and govern many hippocampal-prefrontal-driven cognitive functions, such as spatial working memory (Jones et al., 2005), navigation (Buzsaki, 2013), memory formation (Tambini et al., 2018) and memory integration (Backus et al., 2016). Moreover, given that mid-frontal cortex is a key region of the default mode network (DMN) and that theta oscillations are inversely related to the activation of the DMN (Scheeringa et al., 2007), we suggest that high amyloid load leads to an hypoactivation of the DMN which ultimately manifests as a mid-frontal theta band hyper-activation at the macroscopic scale. DMN functional alteration is reported in a large amount of brain imaging studies on pre-clinical AD individuals with or without cortical amyloid plaques (e.g., Jones et al., 2011; Sheline et al., 2010; Greicius et al., 2009; 2004), and in a recent fMRI study on the INSIGHT-preAD cohort (Chiesa et al., 2019). From a neuro-cognitive point of view, mid-frontal cortices increase of theta rhythmogenesis in response to high amyloid deposition may either suggest a “compensatory brain mechanisms” (i.e., cognitive reserve; Stern, 2012; 2006; 2002) to cope with amyloidosis, or a functional alteration of the DMN. In view of the absence of correlation between mid-frontal theta power and the individuals’ cognitive scores (above all, memory), we tend to suggest that if any compensatory mechanism exists, this may not manifest at a cognitive level, but rather at a mere functional one. It may however be noted that our data might not sufficiently inform this debate and the absence of correlation between cognition and physiology could be expected in the present study, because i) the INSIGHT-preAD individuals have been specifically selected for being cognitively intact from an objective point of view, and ii) importantly, no statistical difference exists at a behavioural level across the individuals of the INSIGHT-preAD cohort.

Surprisingly we did not find any effect of the *APOE* genotype on the mid-frontal theta dynamics while theta dynamics was correlated to amyloid load and amyloid load is known to be correlated with the *APOE* genotype (Dubois, 2016; Morris et al., 2010), and pre-symptomatic carriers of the *APOE* ε4 allele with a higher amyloid load show a more rapid cognitive decline as compared to *APOE* ε4 non-carriers (Mormino et al., 2014). However, to the best of our knowledge, only a few structural (Dowell et al., 2016) or functional (Chiesa et al., 2019) neuroimaging studies, as well as a limited number of EEG investigations (Hatz et al., 2013; Canuet et al., 2012) have fed in this hypothesis. In the present work, we can only provide evidence on the predominance of brain amyloidosis over *APOE* on the sources of brain rhythms and suggest further *ad-hoc* investigation to carefully disentangle this issue. Alternatively, *APOE* genotype-independent changes in EEG could be related to tau pathology which has not been assessed in the INSIGHT cohort yet. Future EEG studies should benefit from the combination of neuroimaging techniques and multimodal biomarker data, for example with novel functional neuroimaging genetics which should bring new evidence in the field (Chiesa et al., 2017).

Crucially, longitudinal source-level analyses provide insightful results to help inform the previous debate of any compensatory mechanisms to cope with cortical amyloidosis, and any implication of the DMN. In fact, while we did not replicate the difference at M24 in mid-frontal theta power between A+ and A-, analyses at M24 show that cortical amyloid status discriminated theta band power differences in the pre-cuneus/posterior-cingulate cortex. Neuroscience studies shows a major implication of this area on the AD spectrum. Particularly, while increased pre-cuneus atrophy has been associated to early-AD (Karas et al., 2007), higher amyloid load in this region characterises a reduced cholinergic activity in overt AD (Stricker et al., 2012; Ikonomovic et al., 2011). From a functional perspective, aberrant activity in this area (Rami et al., 2012), as well as a decreased connectivity between the pre-cuneus and the anterior DMN differentiates normal aging from AD (Klaassens et al., 2017) and can even target memory performances in prodromal AD (Koch et al., 2018). Our results are not only in line with these findings, but also highlight for the first time a direct association of cortical amyloid load with the theta-band hyper-activation of the pre-cuneus in individuals at-risk for AD. Importantly, given the pivotal role that the pre-cuneus plays in the DMN (Fransson et al., 2008) and the inverse relation between theta increase and DMN activation (Sheeringa et al., 2007), such an increase of posterior theta oscillations can corroborate the hypothesis of an amyloid-driven hypo-activation of this network. Remarkably, here we provide the first evidence that theta oscillations behave like a non-static neurodynamic in the AD continuum (Muller et al., 2019; Lubenov et al., 2009; Fransson et al., 2008) and that amyloid status modulates theta changes over two pivotal hubs of the DMN, namely the mid-frontal cortex and the pre-cuneus.

Finally, it is worth mentioning the absence of any significant difference in the alpha band, a frequency that has been convincingly associated to AD progression. From a qualitative view point, here we only report a trend to a decrease of alpha band power as a function of the amyloid status at the second-year follow-up, thus suggesting that posterior alpha slowing - which is canonically associated to physiological aging, MCI and AD - may emerge at later stages of the pre-AD. Alternatively, it might be hypothesized that in pre-AD the amyloid-driven alpha power changes also occur on different brain networks. While in this work we did not confirm this result, in a recent EEG study from the INSIGHT cohort (data not shown) we noticed an increase of alpha band activity in the frontal scalp-electrodes associated with an increased amyloid uptake, which raises a plateau at the threshold of amyloid positivity and then diminishes as amyloid burden increases.

## 5. Conclusions

In conclusion, we provide the first longitudinal evidence on the impact of brain amyloidosis on the EEG dynamics of a large-scale, monocentric AD cohort. We show that different neural markers are in play at different time points of the follow-up. Theta band power increase seems to play a crucial role in pre-AD. Moreover, theta power highlights a potential travelling property, such that its plastic transition from the anterior to the posterior DMN characterised the two stages of the follow-up. Noteworthy, alpha band power slowing showed a decremental tendency at the second-year follow-up, albeit with a relatively poor statistical power. Future follow-up studies on the INSIGHT-preAD cohort will allow further tracking of the evolution of such EEG dynamics.

## Funding

The INSIGHT-preAD study was promoted by INSERM in collaboration with ICM, IHU-A-ICM, and Pfizer and has received a support within the “Investissement d’Avenir (ANR-10-AIHU-06 and ANR-11-INBS-0006). This research publication benefits from the support of the “Big Brain Theory” Program of the ICM (LIBERATE project).

## Conflict of Interest statement

The authors declare no conflict of interest.

## Acknowledgments

The INSIGHT-preAD Study Group:

Audrain C, Auffret A, Bakardjian H, Baldacci F, Batrancourt B, Benakki I, Benali H, Bertin H, Bertrand A, Boukadida L, Cacciamani F, Causse V, Cavedo E, Cherif Touil S, Chiesa PA, Colliot O, Dalla Barba G, Depaulis M, Dos Santos A, Dubois B, Dubois M, Epelbaum S, Fontaine B, Francisque H, Gagliardi G, Genin A, Genthon R, Glasman P, Gombert F, Habert MO, Hampel H, Hewa H, Houot M, Jungalee N, Kas A, Kilani M, La Corte V, Le Roy F, Lehericy S, Letondor C, Levy M, Lista S, Lowrey M, Ly J, Makiese O, Masetti I, Mendes A, Metzinger C, Michon A, Mochel F, Nait Arab R, Nyasse F, Perrin C, Poirier F, Poisson C, Potier MC, Ratovohery S, Revillon M, Rojkova K, Santos-Andrade K, Schindler R, Servera MC, Seux L, Simon V, Skovronsky D, Thiebaut M, Uspenskaya O, Vlaincu M.

## Abbreviations

AD: Alzheimer’s disease
DMN: default mode network
EEG: electroencephalogram
(f)MRI: (functional) magnetic resonance imaging
MCI: Mild Cognitive Impairments
PET: positron emission tomography
pre-AD: pre-clinical Alzheimer’s disease

## Notes

### Competing Interest Statement

The authors have declared no competing interest.

